# mTORC1 in AGRP neurons integrates exteroceptive and interoceptive food-related cues in the modulation of adaptive energy expenditure in mice

**DOI:** 10.1101/110544

**Authors:** Luke K Burke, Tamana Darwish, Althea R Cavanaugh, Sam Virtue, Emma Roth, Joanna Moro, Shun-Mei Liu, Jing Xia, Jeffrey W. Dalley, Keith Burling, Streamson Chua, Toni Vidal-Puig, Gary J Schwartz, Clémence Blouet

## Abstract

Energy dissipation through interscapular brown adipose tissue (iBAT) thermogenesis is an important contributor to adaptive energy expenditure. However, it remains unresolved how acute and chronic changes in energy availability are detected by the brain to adjust iBAT activity and maintain energy homeostasis. Here we provide evidence that AGRP inhibitory tone to iBAT represents an energy-sparing circuit that integrates environmental food cues and internal signals of energy availability. We establish a role for the nutrient-sensing mTORC1 signaling pathway within AGRP neurons in the detection of environmental food cues and internal signals of energy availability, and in the bi-directional control of iBAT thermogenesis during nutrient deficiency and excess. Collectively, our findings provide insights into how mTORC1 signaling within AGRP neurons surveys energy availability to engage iBAT thermogenesis, and identify AGRP neurons as a neuronal substrate for the coordination of energy intake and adaptive expenditure under varying physiological and environmental contexts.

---

In mammals, the maintenance of energy balance relies on a tight coordination of energy intake and energy expenditure. Internal energy surfeit, typically achieved via maintenance on energy dense food in a laboratory setting, promotes recruitment of interscapular brown adipose tissue (iBAT) thermogenesis, increasing energy expenditure and therefore limiting weight gain^1,2^. Conversely, energy expenditure is robustly decreased in energy-restricted rodents and humans^3,4^. This protective mechanism mitigates weight loss when environmental energy sources are limited and may contribute to the failure of long-term maintenance of weight loss through dieting or pharmacological interventions. The development of successful therapies to treat obesity therefore requires a better characterization of the mechanisms underlying the regulation of adaptive energy expenditure in various contexts of energy availability.

Sympathetic nervous outflow to iBAT modulates adaptive energy expenditure by promoting iBAT thermogenesis ^5^ and is modulated by internal energy availability ^6^. Melanocortinergic circuits, one of the best characterized central energy-sensing network regulating energy balance, have been implicated in the control of sympathetic outflow to iBAT ^5,7^, but the exact neuronal mechanisms and energy-sensing pathways that modulate sympathetic tone to iBAT during caloric restriction or nutrient excess have not been identified.

Orexigenic neurons expressing agouti-related peptide (AGRP) and neuropeptide Y (NPY) form a discrete population of about 10,000 cells residing in the arcuate nucleus of the mouse hypothalamus (ARH). These neurons respond to bidirectional changes in energy availability, are activated by fasting, low glucose levels and ghrelin, and are inhibited by leptin, insulin, refeeding and high glucose levels ^8-9^. Their essential role in the bidirectional control of energy intake can be appreciated by the aphagia observed in mice with specific ablation of this neuronal population during adulthood ^10,11^, and the voracious feeding response upon opto- or chemogenetic activation of AGRP neurons in sated mice ^12,13^. Initial evidence indicates that AGRP neurons also regulate energy expenditure ^13^, but the contribution of iBAT thermogenesis to this effect, the physiological contexts in which this axis is engaged, and the mechanisms through which AGRP neurons monitor energy availability have not been identified.

Here we used chemogenetics to rapidly and selectively activate AGRP neurons, and demonstrate that AGRP neurons can engage an energy-sparing circuit to iBAT that represents the integration of metabolic interoception and environmental food-related cues. We show that mTORC1 signaling in AGRP neurons surveys sensory food cues and internal signals of energy availability. Using inducible lentivectors to manipulate the nutrient-sensing mTORC1 signaling pathway specifically in AGRP neurons of adult normally developed mice, we show that mTORC1 signaling within AGRP neurons acts as a bidirectional node of control that modulates the AGRP-iBAT axis. Collectively, our data describe a novel neuronal energy sensing mechanism that modulates sympathetic tone to iBAT and identify AGRP neurons as a neuronal substrate for the coordination of energy intake and adaptive expenditure under various physiological and environmental contexts.

## RESULTS

### Activation of AGRP neurons rapidly inhibits sympathetic tone to iBAT and suppresses iBAT thermogenesis

To rapidly and reversibly activate AGRP neurons, we used the stimulatory hM3dq designer receptor exclusively activated by designer drugs (DREADD) ^14^. This receptor couples through the Gq pathway upon exposure to the otherwise pharmacologically inert ligand clozapine-N-oxide (CNO), leading to neuronal depolarization. *Agrp-IRES-cre* mice received a bilateral stereotactic injection of an adeno-associated virus expressing hM3dq (AAV-hM3dq-mCherry), and were implanted with a telemetric probe under the interscapular adipose depot to remotely monitor iBAT temperature in behaving animals. Successful stereotactic targeting of hM3dq in AGRP neurons was confirmed by ARH mCherry staining (Figure 1 – Figure Supplement 1a), as well as with a functional feeding test. Mice selected for subsequent studies ate at least 2-fold more during the first hour following the CNO injection as compared to the vehicle injection (Figure 1 – Figure Supplement 1b).

**Figure 1.**
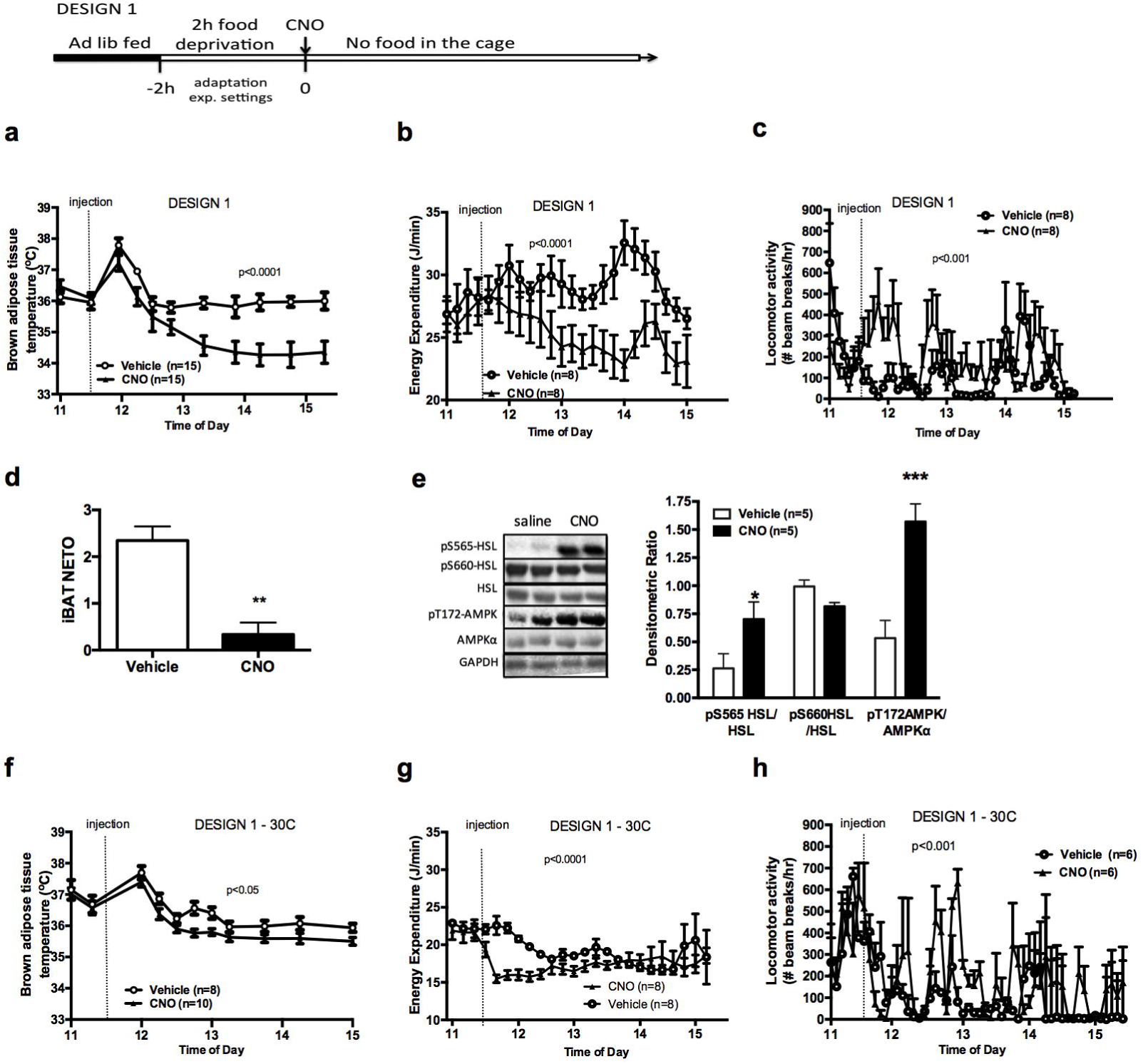
AGRP neurons control iBAT thermogenesis. iBAT temperature (a), energy expenditure (b), locomotor activity (c), Norepinephrine turnover (n=8) (d) and phosphorylation levels of lipolytic pathways in the iBAT (d) following chemogenetic activation of AGRP neurons in food-deprived mice (DESIGN 1). iBAT temperature (f), energy expenditure (g) and locomotor activity (h) following chemogenetic activation of AGRP neurons at 30°C. Data are mean ± SEM, *: p<0.05; ***: p<0.001.

Previous reports indicate that chemogenetic activation of AGRP neurons rapidly increases food intake and suppresses oxygen consumption ^13^. To determine the direct effect of AGRP activation on adaptive energy expenditure independently of energy intake, we tested the effect of AGRP activation on iBAT temperature and energy expenditure in mice that were food deprived for 2h, followed by an ip injection of CNO or vehicle without access to food following the injection (Design 1). As expected, activation of AGRP neurons rapidly suppressed iBAT temperature (Fig. 1a) and energy expenditure (Fig. 1b) despite increasing locomotor activity (Fig. 1c) when mice were maintained in a food-free environment following CNO administration. Local administration of ghrelin, an endogenous activator of AGRP neurons, into the mediobasal hypothalamus (MBH) of wild-type mice decreased iBAT temperature (Figure 1 – Figure Supplement 1c), supporting the physiological relevance of this effect.

We investigated the mechanisms mediating the rapid fall in iBAT thermogenic activity following AGRP neuronal activation by first assessing sympathetic outflow to iBAT. iBAT norepinephrine turnover (NETO) was measured using the α-methyl-p-tyrosine (AMPT) method. In CNO-treated mice, iBAT NETO was significantly lower than in vehicle-treated mice (Fig. 1d), supporting the conclusion that AGRP activation rapidly suppresses sympathetic output to iBAT. In contrast, circulating thyroxine (T4) and FGF-21 levels remained unchanged during the first 45 min following CNO administration (Figure 1 – Figure Supplement 1d and 1e), ruling out the contribution of these hormones in the rapid decrease in iBAT thermogenesis following AGRP neuronal activation. Consistent with previous reports ^15^, we did not observe any changes in iBAT expression of thermogenic genes within an acute 1h time-frame following CNO administration (Figure 1 – Figure Supplement 1f). These results indicate that iBAT recruitment is not involved in the rapid decrease in iBAT temperature observed as soon as 1h after AGRP neuronal activation.

We examined the activation status of signaling pathways downstream from β3-adrenergic receptor in iBAT 1h after CNO administration. We did not detect changes in the expression or activation status of PKA, CREB, perilipin, UCP-1 or in the activating phosphorylation of HSL on Ser^660^ (Figure 1 – Figure Supplement 1g). In contrast, inhibitory phosphorylation of HSL on Ser^565^ was significantly increased in the iBAT of CNO-treated mice (Fig. 1e). AMP-activated protein kinase (AMPK) is inhibited by β3-adrenergic signaling in iBAT and has been reported to phosphorylate HSL on Ser^565^ phosphorylation ^16,17^. This led us to hypothesize that decreased sympathetic tone to iBAT in response to AGRP activation may activate AMPK and reduce iBAT thermogenesis via an increase in HSL inhibitory phosphorylation on Ser^565^. Accordingly, we observed a robust increase in AMPK activatory phosphorylation in the iBAT of CNO-treated mice (Fig. 1e).

To determine whether the control of iBAT thermogenesis by AGRP neurons was independent of thermoregulation, we repeated these experiments in mice maintained in the thermoneutral zone (30°C), a condition under which thermoregulatory thermogenesis is abolished. Under these conditions, the effects of AGRP activation on iBAT temperature, energy expenditure and locomotor activity were attenuated, but remained statistically significant (Fig. 1f-1h). This result further directly implicates iBAT thermogenic activity in the response to AGRP activation and indicates that the AGRP-iBAT circuit can be engaged independently of thermoregulation. At 4°C, a condition under which thermoregulatory iBAT thermogenesis is strongly activated ^6^, AGRP neuronal activation reduced iBAT temperature to a similar extent as at ambient temperature (Figure 1 – Figure Supplement 1h), reinforcing the idea that the dynamic range of AGRP-iBAT circuit function is dissociable from environmental temperature and cold-induced thermoregulation. Thus, activation of AGRP neurons increases locomotor activity and produces a substantial sustained decrease in energy expenditure through a suppression of iBAT thermogenic activity.

### The regulation of iBAT thermogenesis by AGRP neurons depends on food availability and is modulated by sensory cues of environmental food availability

To better characterize the relationship between the control of energy intake and energy expenditure in response to AGRP neuronal activation, we performed experiments in mice that had access to food following CNO administration (Design 2). Under these conditions, activation of AGRP neurons increased locomotor activity and energy intake but failed to alter iBAT temperature and energy expenditure (Fig. 2a-2c), indicating that energy availability modulates the AGRP-iBAT circuit.

**Figure 2.**
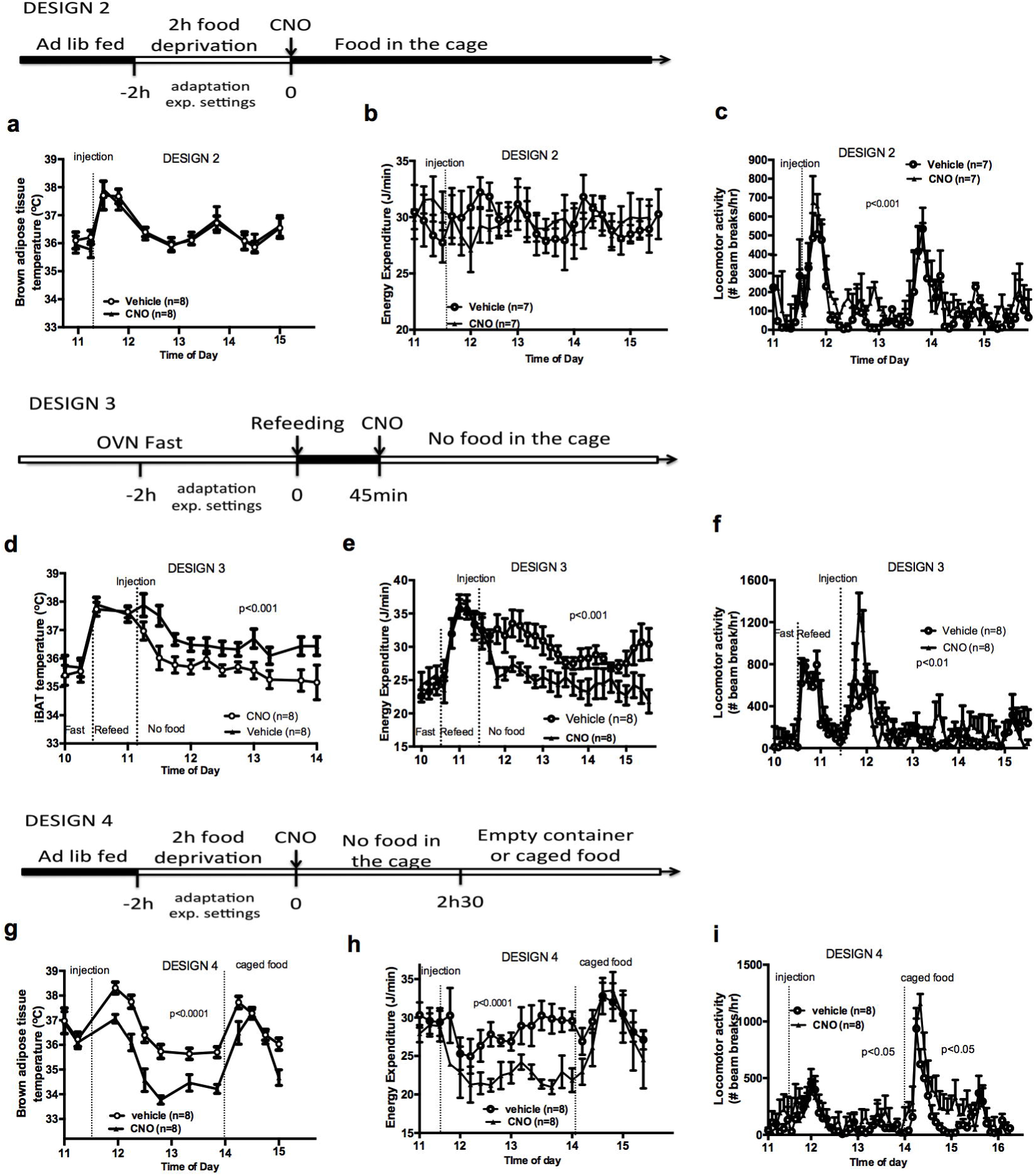
Food availability and sensory cues of food availability modulate the AGRP-iBAT circuit. iBAT temperature (a), energy expenditure (b) and locomotor activity (c) following chemogenetic activation of AGRP neurons in the presence of food (DESIGN 2). iBAT temperature (d), energy expenditure (e) and locomotor activity (f) following chemogenetic activation of AGRP neurons in refed mice (DESIGN 3). iBAT temperature (g), energy expenditure (h) and locomotor activity (i) following chemogenetic activation of AGRP neurons in the presence of caged food (DESIGN 4). Data are mean ± SEM, *: p<0.05; ***: p<0.001.

To test whether internal signals of energy availability were sufficient to blunt the response to CNO, mice were fasted overnight and refed before receiving an injection of CNO or vehicle, and then placed in food-free clean cages immediately after the injection (Design 3). In that context, AGRP neuronal activation suppressed iBAT temperature and energy expenditure (Fig. 2d-2f), suggesting that internal signals of energy availability are not sufficient to blunt the AGRP-iBAT hypometabolic response.

We noticed that iBAT temperature and energy expenditure were rapidly restored in CNO-treated mice upon food reintroduction in design 1 (Figure 2 – Figure Supplement 1a and 1b), suggesting that the presence of food rapidly resets the AGRP-iBAT circuit. To directly test the role of environmental cues of food availability in AGRP inhibitory tone to iBAT, we repeated this experiment in mice kept in clean food-free cages after CNO administration and transiently exposed to food sensory stimuli. Specifically, we introduced chow pellets in a container that allowed food to be seen and smelled but not consumed (caged-food). This paradigm was recently found to rapidly inhibit AGRP neurons in hungry mice ^8^. Presenting caged-food to CNO-treated mice rapidly increased iBAT temperature (Fig. 2g) and energy expenditure (Fig. 2h) to levels similar to those measured in controls, an effect that was transient and lasted for about 1h. Importantly, presentation of an inedible empty container to naïve animals failed to alter iBAT temperature and energy expenditure (Figure 2 – Figure Supplement 1c-1d), indicating that the stress response to the introduction of an object to naïve mice is not sufficient to produce the restoration in iBAT temperature seen with presentation of food sensory cues. Again we observed that CNO administration increased locomotor activity (Fig. 2i) despite reducing energy expenditure and iBAT temperature. These results indicate that pre-ingestive food detection is sufficient to apprehensively turn off the AGRP-iBAT circuit suppressing iBAT thermogenesis and energy expenditure. Thus, the AGRP-iBAT circuit integrates sensory and endogenous signals of energy availability in the regulation of iBAT thermogenic activity and energy expenditure to promote energy sparing when fuel availability is scarce.

### mTORC1 signaling in AGRP neurons responds to acute and chronic signals of energy abundance and environmental cues of food availability

AGRP neurons are essential to the control of energy balance and ideally positioned within and around the median eminence to detect circulating signals of energy availability. However, little is known about the intracellular energy-sensing signaling pathways coupling energy availability to neuronal activity in AGRP neurons. The energy-sensing mTORC1 pathway is an evolutionary-conserved neuronal nutritional and hormonal sensor that regulates hunger-driven eating and foraging ^18-19^, two behaviors regulated by AGRP neurons ^12,13,20^. We hypothesized that energy sensing through mTORC1 in AGRP neurons monitors internal energy availability to coordinate adaptive energy expenditure.

We first tested whether the activity of mTOR in NPY/AGRP neurons is modulated by a refeeding episode. This nutritional transition is associated with a rapid inhibition of AGRP neurons ^8^, an activation of iBAT thermogenic activity and an increase in energy expenditure (Figure 3 – Figure Supplement 1a and 1b). We used immunofluorescence to detect the active form of mTOR phosphorylated at Ser^2448^ (p-mTOR) in hypothalamic slices. Rapamycin suppressed p-mTOR expression (Figure 3 – Figure Supplement 1c), confirming the specificity of this staining. Refeeding for 1h following an overnight fast significantly increased p-mTOR expression in ARH NPY neurons (Fig. 3a and 3d). Importantly, p-mTOR activation following refeeding occurs only in 15% of ARH NPY/AGRP neurons, supporting the conclusion that postprandial changes in energy availability is coupled to changes in mTOR signaling only in a subpopulation of ARH NPY/AGRP neurons. Costained neurons were mostly absent from the rostral ARH (−1.22 to −1.6mm post Bregma ^21^) and concentrated in the ventro-medial neurons of the caudal ARH (−1.7mm to −2mm post bregma ^21^) (Figure 3 – Figure Supplement 1d).

**Figure 3.**
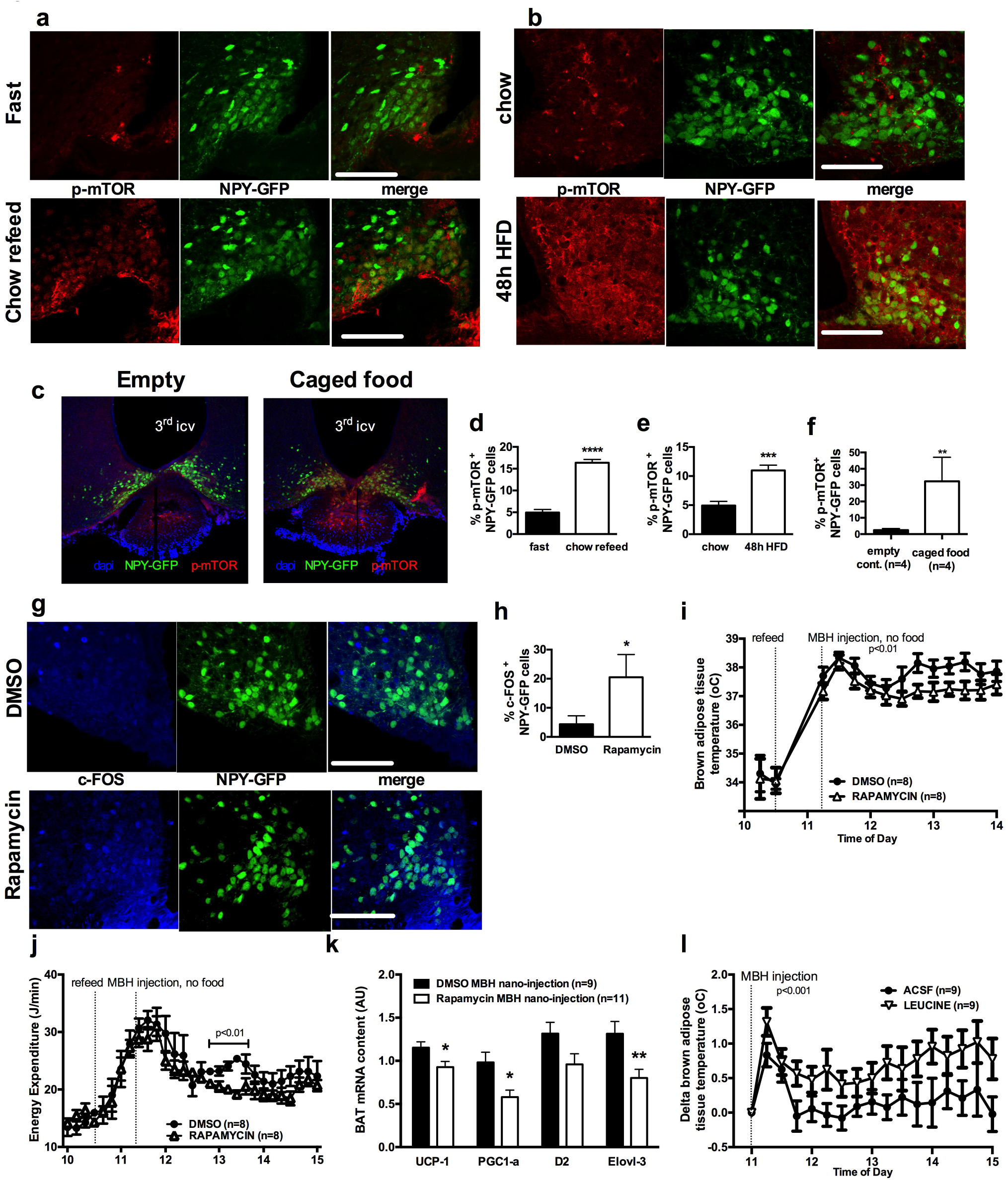
mTOR signaling in AGRP neurons senses endogenous and environmental signals of energy availability, regulates AGRP activity and iBAT thermogenesis. Activation levels of mTOR in NPY-GFP neurons following a 18h fast and 1h chow re-feed (a, d, n=5), and a 48h transition from chow to HF diet (b, e, n=6), and a 18h fast followed by a 30 min exposure to caged food (c, f). NPYGFP
neuronal c-fos expression following inhibition of mTORC1 using rapamycin (g, h) – scale bar is 100μm. iBAT temperature (i), energy expenditure (j) and iBAT thermogenic gene expression (k) following MBH nanoinjection of rapamycin in refed mice. iBAT temperature (l) following MBH nanoinjection of L- leucine in fasted mice. Data are mean ± SEM *: p<0.05; ***: p<0.001; ****: p<0.0001.

Another metabolic context associated with a rapid activation of iBAT thermogenesis is exposure to a high fat (HF) diet ^22^. We confirmed that after 2 days of HF feeding, iBAT thermogenesis was increased (Figure 3 – Figure Supplement 1e and 1f) leading to an increase in energy expenditure (Figure 3 – Figure Supplement 1g), as previously reported ^23^. We observed that mTOR signaling was also upregulated in NPY/AGRP neurons in this context, again only in a small group of this neuronal population (Fig. 3b and 3e). These data indicate that in nutritional conditions under which iBAT thermogenic activity is engaged, mTOR activity is increased in a subpopulation of AGRP/NPY neurons in which metabolic signals are coupled to changes in mTOR signaling. These findings raise the possibility that mTOR in AGRP neurons mediate changes in sympathetic tone to iBAT under these conditions.

Given our observations that the inhibition of iBAT thermogenesis induced by AGRP neuronal activation is abolished in the presence of food, we sought to assess whether hypothalamic mTOR signaling is also sensitive to food sensory stimuli. We first measured p-mTOR expression in NPY/AGRP neurons of overnight fasted mice exposed for 30 min to an empty contained or caged food. In control mice exposed to an empty container, p-mTOR was absent from ARH NPY-GFP neurons (Figure 3c and 3f). In contrast, 30% ARH NPY-GFP neurons expressed p-mTOR in response to 30min exposure to caged food. These data indicate that mTOR signaling in AGRP neurons, in addition to responding to acute (refeeding) and chronic (48h HF feeding) internal signals of energy availability, also responds to environmental food-related cues.

### MBH mTORC1 signaling modulates AGRP neuronal activity and iBAT thermogenesis

To directly probe the link between mTOR activity and AGRP neuronal activity, we treated refed *NPY-GFP* mice with rapamycin, a well-characterized mTORC1 inhibitor, and quantified the expression of the marker of neuronal activation c-fos in NPY-GFP cells. Rapamycin increased the expression of cfos in NPY-GFP neurons (Fig. 3g and 3h), supporting the interpretation that mTORC1 signaling negatively regulates the activity of AGRP/NPY neurons.

We then tested whether acute changes in mTORC1 activity in AGRP neurons modulate iBAT thermogenesis using nanoinjections of activators and inhibitors of the mTORC1 pathway into the MBH of wild-type mice. Rapamycin diluted in 100% DMSO was administered into the hypothalamic parenchyma in nanovolumes to acutely activate mTORC1 signaling. Rapamycin-mediated inhibition of MBH mTORC1 in refed mice significantly reduced iBAT temperature (Fig. 3i), energy expenditure (Fig. 3j), and altered iBAT thermogenic gene expression (Fig. 3k) without significantly affecting locomotor activity (Figure 3 – Figure Supplement 1h). Of note, although this DMSO concentration may induce cellular stress responses, the low volume bolus we injected did not produce noticeable changes in behavior, locomotor activity, energy intake and energy expenditure compared to non-injected mice under the same conditions (not shown). Conversely, activation of MBH mTORC1 using the amino acid L-leucine produced an increase in iBAT temperature in fasted mice (Fig. 3l) but did not produce changes in energy expenditure (Figure 3 – Figure Supplement 1i). Thus, acute changes in MBH mTORC1 signaling are associated with bidirectional changes in iBAT thermogenesis.

### Constitutive activation of mTORC1 signaling in AGRP neurons increases iBAT thermogenesis and energy expenditure and protects against diet-induced obesity

To directly test the role of increased mTORC1 signaling specifically in AGRP neurons in the control of iBAT thermogenesis, we generated a lentivector expressing a constitutively active mutant of the mTORC1 effector p70 S6 kinase 1 (S6K1) in a cre-dependent manner (pCDH-CMV-FLEX-HA-S6K1-F5a). We delivered the pCDH-CMV-FLEX-HA-S6K1-F5a vector into the ARH of normally-developed adult *Agrp-IRES-cre* or WT mice (mice subsequently referred to as Agrp-CA-S6 and WT-CA-S6) using stereotactic nanoinjections. *Agrp-IRES-cre;Npy-gfp* were used to validate the construct. Ha staining indicated that over 95% of the cells expressing the construct were NPY-GFP neurons and that over 60% of NPY-GFP cells expressed the construct (Figure 4 – Figure Supplement 1a). Increased expression of the active form of ribosomal protein S6 (phosphorylated at Ser 240-244, p-rpS6, main downstream effector of S6K) in ARH NPY-GFP neurons confirmed constitutive activation of the pathway in AGRP neurons of Agrp-CA-S6 mice compared to controls (Figure 4 – Figure Supplement 1b).

**Figure 4.**
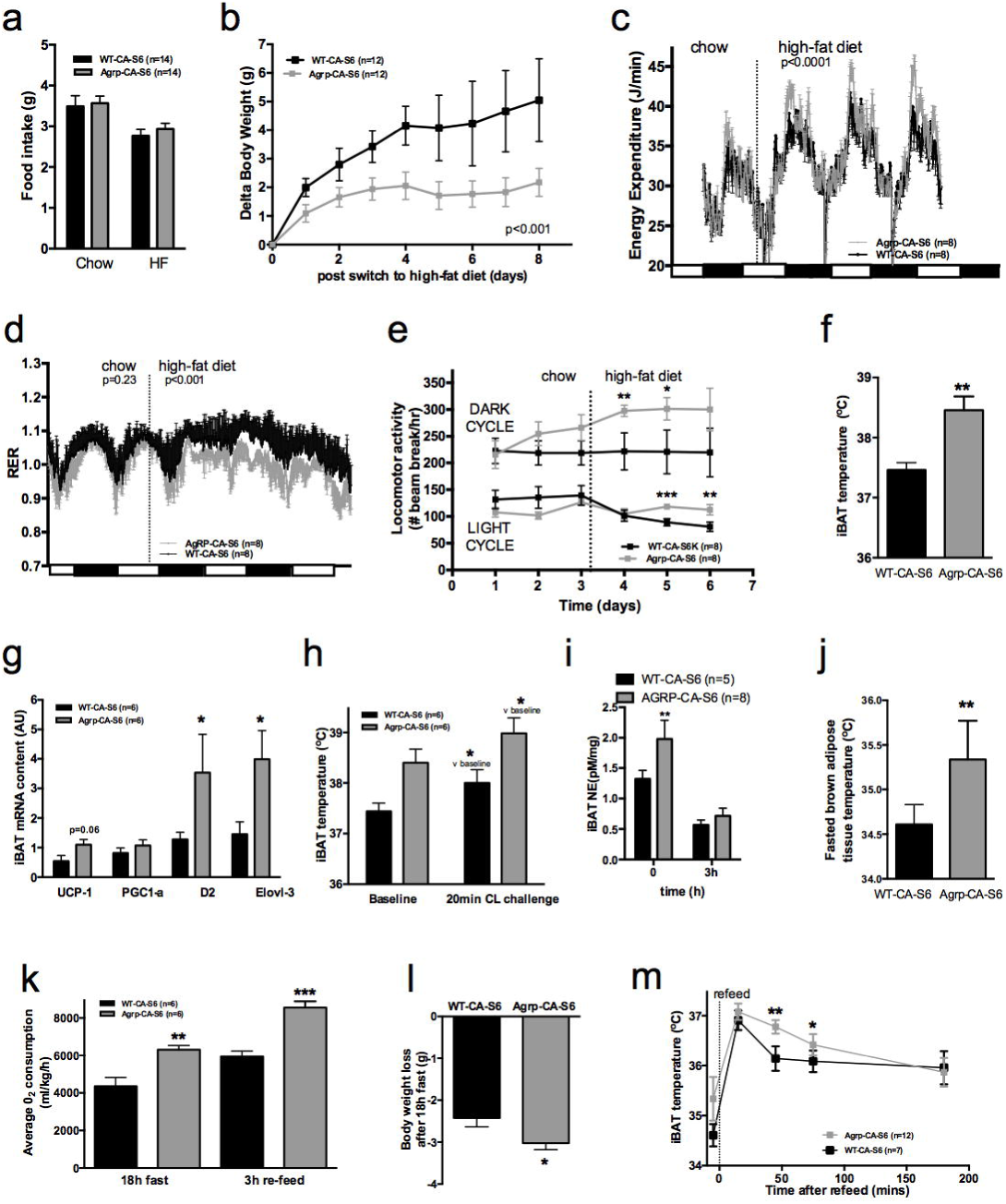
Metabolic phenotyping of mice expressing constitutively active mutant of mTORC1 effector p70 S6 kinase 1 in *Agrp-IRES-cre* (Agrp-CA-S6) or wild-type (WT-CA-S6) littermates. Food intake (a), body weight gain on high-fat diet (b), energy expenditure (c), RER (d), locomotor activity (e), iBAT temperature (f), iBAT thermogenic gene expression (g), iBAT temperature following β3- agonist-induced challenge (h), and Norepinephrine turnover (i) in Agrp-CA-S6 and WT-CA-S6 mice. iBAT temperature (i), energy expenditure (j), and body weight loss during 18h overnight fast (k), and iBAT temperature during 3h re-feed (l) in Agrp-CA-S6 and WT-CA-S6 mice. Data are mean ± SEM *: p<0.05; **: p<0.01; ***: p<0.001.

Constitutive activation of S6K1 in AGRP neurons had no effect on food intake (Fig. 4a), body weight gain (Figure 4 – Figure Supplement 1c), adiposity (Figure 4 – Figure Supplement 1d) or glucose tolerance (Figure 4 – Figure Supplement 1e) when mice were maintained on a chow diet. Given the crucial role of AGRP neurons in feeding behavior and survival ^10^, these results indicate that manipulation of mTORC1 signaling in AGRP neurons does not produce loss of AGRP function.

Upon exposure to a HF diet, Agrp-CA-S6 mice were protected against body weight gain (Fig. 4b). We performed a detailed characterization of energy balance during the transition from chow to HF diet. Constitutive activation of S6K1 in AGRP neurons did not affect energy intake (Fig. 4a) but produced a significant increase in night-time energy expenditure under HF diet (Fig. 4c). The specific implication of energy expenditure in this metabolic phenotype, and the preservation of feeding behavior, support the conclusion that chronic changes in S6K1 signaling in AGRP neurons do not produce an overall change in AGRP activity, but rather modulate the activity of a specific subpopulation of AGRP neurons regulating energy expenditure. Substrate utilization was similar between groups on chow but fat utilization was substantially increased after the switch to the HF diet in Agrp-CA-S6 mice (Fig. 4d). Likewise, locomotor activity significantly increased in Agrp-CA-S6 mice after the transition to the HF diet (Fig. 4e). Night-time iBAT temperature was significantly increased in Agrp-CA-S6 mice (Fig. 4f) and consistently, mRNA expression of iBAT thermogenic markers was increased (Fig. 4g), suggesting a role for increased iBAT activation in the elevated energy expenditure and reduced weight gain observed in these mice. The amplitude of iBAT thermogenic response to β-adrenergic stimulation was similar between groups (Fig. 4h). This indicates that elevated iBAT thermogenesis in *Agrp-IRES-cre* mice was not the result of an impairment in iBAT responsiveness to sympathetic activation. Instead, increased iBAT norepinephrine turnover in AGRP-CA-S6 mice compared to control indicated increased sympathetic tone to iBAT following constitutive activation of S6K in AGRP neurons (Fig. 4i; NETO: −0.75 pM/mg/h vs. −1.26 pM/mg/h in WT-CA-S6 and AGRP-CA-S6 respectively, p<0.05). Constitutive activation of S6K1 in AGRP neurons also increased iBAT temperature (Fig. 4j) energy expenditure (Fig. 4k) during a fast resulting in enhanced body weight loss (Fig. 4l). Postprandial increases in iBAT thermogenesis and energy expenditure were also higher in mice expressing constitutively active S6K1 in AGRP neurons (Fig. 4l, 4m). Thus, constitutive activation of S6K signaling in AGRP neurons of adult normally developed mice protect against diet-induced weight gain through an increase in sympathetic tone to iBAT and an increase in iBAT thermogenesis.

### Inhibition of endogenous mTORC1 in AGRP neurons reduces iBAT thermogenesis and energy expenditure and promotes HF-induced weight gain

In order to investigate the role of endogenous mTORC1 signaling in AGRP neuronal control of energy homeostasis, we generated a lentivector expressing a kinase dead mutant of the S6K1 in a cre-dependent manner (pCDH-CMV-FLEX-HA-S6K1-KR) and injected this virus into the MBH of normally developed adult *Agrp-IRES-cre* mice or wild-type littermates through local stereotatic nanoinjections (Agrp-DN-S6 and WT-DN-S6). *Agrp-IRES-cre;Npy-gfp* were used to validate the construct. Ha staining indicated that over 95% of the cells expressing the construct were NPY-GFP neurons and that over 40% of NPY-GFP cells expressed the construct (Figure 5 – Figure Supplement 1a). Decreased expression of p-rpS6 in ARH NPY-GFP neurons confirmed inhibition of the pathway in AGRP neurons of Agrp-DN-S6 mice compared to controls (Figure 5 – Figure Supplement 1b).

**Figure 5.**
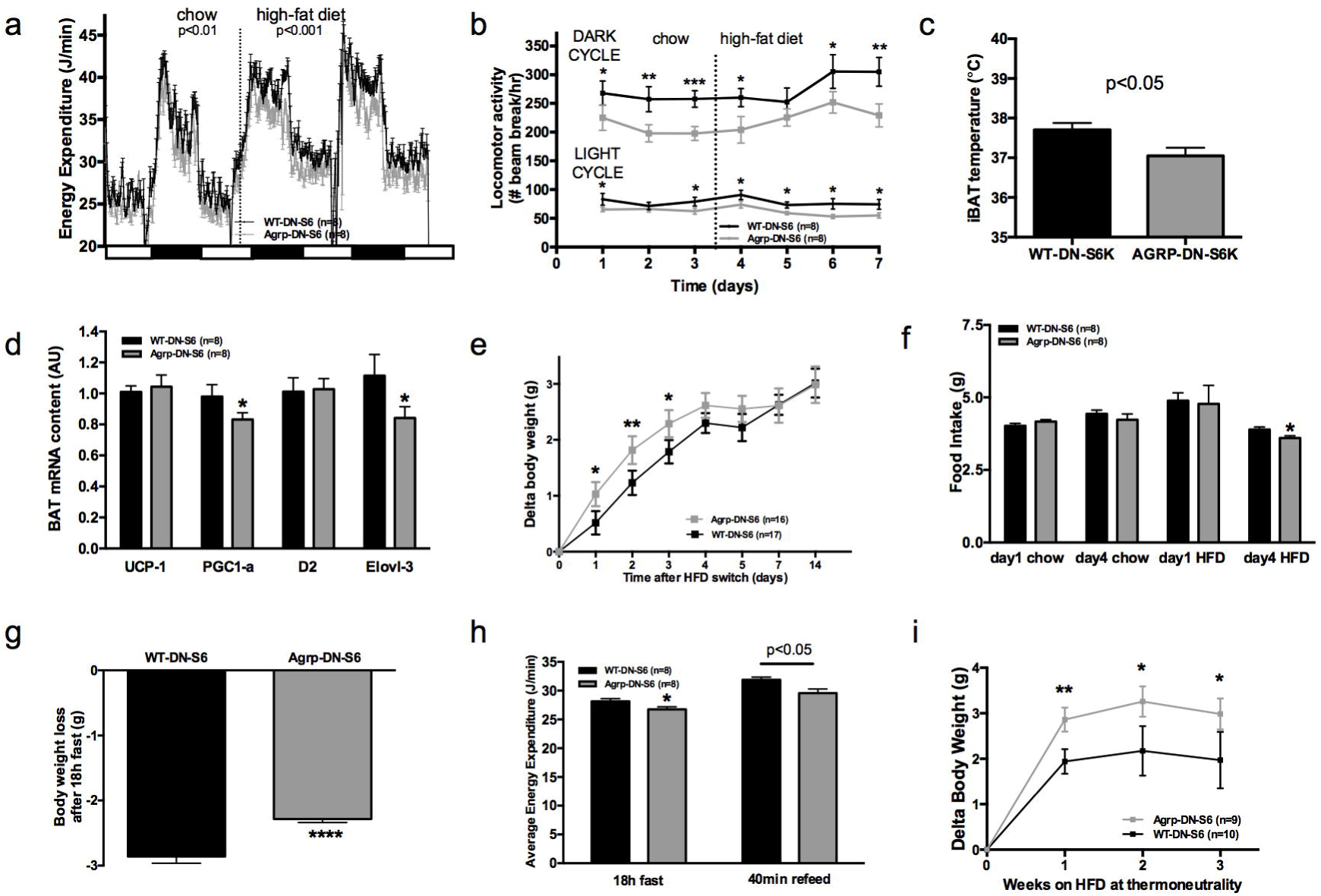
Metabolic phenotyping of mice expressing dominant negative mutant of mTORC1 effector p70 S6 kinase 1 (S6K1) in *Agrp-IRES-cre* (Agrp-DN-S6) or wild-type (WT-DN-S6). Energy expenditure (a), locomotor activity (b), iBAT temperature (c), iBAT thermogenic gene expression (d), body weight gain during initial switch to HF diet (e), food intake (f), body weight loss during overnight 18h fast (g), and energy expenditure during fast and re-feed (h) in Agrp-DN-S6 and WT-DN-S6 mice. Body weight gain of Agrp-DN-S6 and WT-DN-S6 mice at thermoneutrality under chow (i) and HF (k) diet. Data are mean ± SEM *: p<0.05; **: p<0.01; ***: p<0.001.

Quenching endogenous S6K1 in AGRP neurons produced changes in energy expenditure reciprocal to those observed following constitutive activation of the pathway. These were not sufficient to affect body weight gain during chow maintenance (Figure 5 – Figure Supplement 1c) but produced a significant decrease in energy expenditure (Fig. 5a), locomotor activity (Fig. 5b), iBAT temperature (Fig. 5c), and iBAT thermogenic gene expression (Fig. 5d) after switch to a HF diet. Downregulation of S6K1 activity in AGRP neurons only produced a modest reduction in weight gain during the initial switch to HFD (Fig. 5e) during which time we did not observe any differences in food intake (Fig. 5f) or RER (Figure 5 – Figure Supplement 1d). However, we observed a reduction in food intake in DN-S6K mice 4 days after the introduction of the HF diet (Fig. 5f), an effect that likely occurred secondary to reduced energy expenditure and mitigated changes in weight gain.

To further characterize energy expenditure without the confound of energy intake, we challenged these mice with an overnight fast. Despite showing no difference in body weight under *ad libitum* access to food, Agrp-DN-S6 mice lose significantly less weight than wild-type controls during an overnight fast (Fig. 5g), indicating increased energy efficiency in these mice. Consistently, energy expenditure was lower in Agrp-DN-S6 mice throughout the fast (Fig. 5h). In addition, postprandial increases in energy expenditure (Fig. 5h) during the subsequent refeed was lower. Last, body weight gain at thermoneutrality was significantly higher in Agrp-DN-S6 mice than in controls (Fig. 5i). Collectively these data indicate that downregulation of S6K signaling in AGRP neurons leads to a decrease in energy expenditure mediated at least in part by a decrease in iBAT thermogenesis, which leads to a significant increase in body weight gain when mice are maintained at thermoneutrality.

### mTORC1 in AGRP neurons mediates the thermogenic effect of leptin

Central leptin engages the sympathetic nervous system to increase iBAT thermogenesis through the activation of distributed leptin-receptor expressing neurons in the DMH, nMPO, NTS and ARH. Several lines of evidence support a role for leptin receptors in the ARH in leptin-induced activation of iBAT ^24,25^. In the adult brain, leptin inhibits the vast majority of AGRP neurons ^26^. However, leptin-induced hyperpolarization of AGRP neurons has not been implicated in leptin-induced increase in sympathetic tone to iBAT. Previous work implicated hypothalamic mTORC1 signaling in leptin’s effects on energy balance ^18,27^. Thus, we hypothesized that leptin activates mTORC1 signaling in AGRP neurons, leading to an increase in sympathetic tone to iBAT.

To test this hypothesis, we first colocalized markers of mTORC1 signaling in NPY/AGRP neurons 30 min following leptin administration. Leptin increased the expression of p-mTOR (Fig. 6a and 6c) and the expression of the active form of one of its effectors, ribosomal protein S6 (p-rpS6, Fig. 6b and 6d) in NPY/AGRP neurons. As expected, leptin increased expression of pSTAT3 in NPY/AGRP neurons, and we observed that 40% of pSTAT-3^+^ NPY/AGRP neurons also expressed the active form of rpS6.

**Figure 6.**
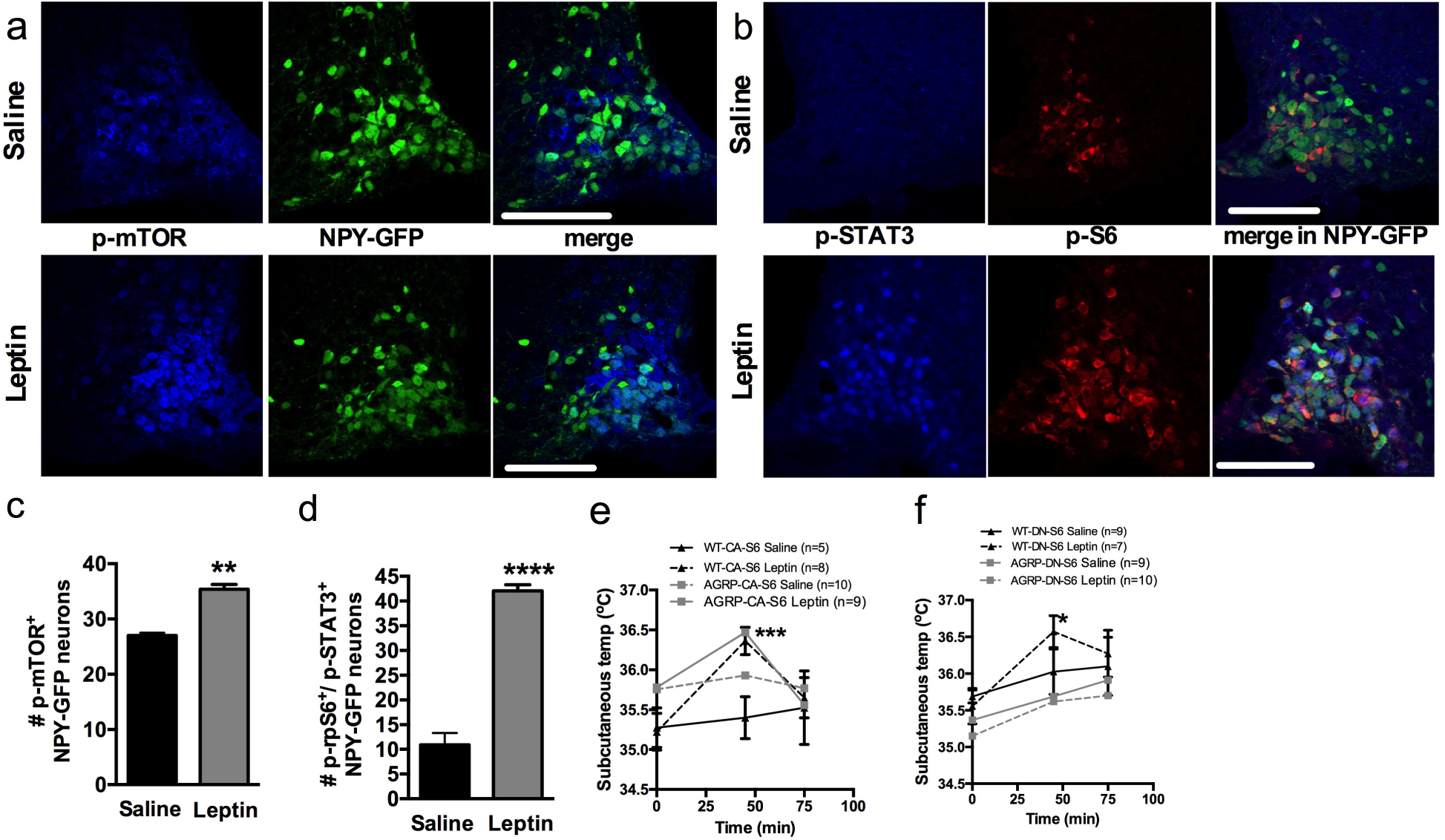
Leptin engages mTORC1 signaling in AGRP neurons to regulate thermogenesis. Expression of p-mTORC1 (a, c) and colocalization of p-rpS6 and p-STAT3 (b, d) in NPY-GFP neurons following systemic leptin treatment. Leptin’s thermogenic effects in WT-CA-S6 and Agrp-CA-S6 (e), and WT-DN-S6 and Agrp-DN-S6 (f) mice. Data are mean ± SEM *: p<0.05; **: p<0.01; ***: p<0.001

We then measured leptin’s thermogenic effect in mice with bidirectional modulations of S6K1 signaling in AGRP neurons. Leptin’s effects on temperature were attenuated in Agrp-CA-S6 mice compared to wild-type controls (Fig. 6e). While both groups reached a similar absolute temperature increase following ip leptin, the relative change in temperature was significantly lower in the Agrp-CA-S6 mice due to their elevated core temperature at baseline. Together these data suggest that lack of flexibility in mTORC1 signaling in AGRP neurons impairs leptin’s thermogenic action. Leptin’s thermogenic effect was completely abolished in Agrp-DN-S6 mice (Fig. 6f), indicating that S6K1 signaling in AGRP neurons is required for leptin’s thermogenic action.

## DISCUSSION

Activation of AGRP neurons has been established as an essential component in the initiation of feeding behavior ^12,13^. Here we demonstrate that the multimodal contribution of AGRP neurons to the maintenance of energy balance includes the regulation of iBAT thermogenesis. We show that activation of AGRP neurons induces a rapid suppression of sympathetic output to iBAT, as previously reported ^28^, leading to a decrease in iBAT thermogenesis and energy expenditure. We show that the AGRP-iBAT circuit is selectively engaged to spare internal energy stores in the absence of food and food sensory cues, supporting the notion that AGRP neurons integrate external and internal signals of energy availability to coordinate energy intake and expenditure in conditions of low energy availability. We provide evidence that this regulation also occurs at thermoneutrality and is therefore potentially relevant to human physiology. We identify the mTORC1 signaling pathway as an intracellular mechanism through which AGRP neurons monitor external and internal energy availability, and demonstrate that mTORC1 signaling modulates AGRP activity, sympathetic tone to iBAT, iBAT thermogenesis and energy expenditure. Last, we show that metabolic sensing via mTORC1 in AGRP neurons is required for the regulation of energy expenditure both during nutritional transitions from fasting to feeding and during adaptation to HF feeding. Our data provide a physiological framework for the role of AGRP neurons in the control of adaptive thermogenesis and the coordination of energy intake and energy expenditure, and represent the first characterization of the functional consequences of AGRP sensory integration in the regulation of energy balance.

A previous study implicated AGRP neurons in the regulation of energy expenditure ^13^. In that study, chemogenetic activation of AGRP neurons produced a rapid reduction in oxygen consumption. Although the effector mediating this decrease had not been investigated, these results are in line with our findings and support a role for AGRP neurons in the regulation of energy expenditure. However, the reduction in oxygen consumption occurred in the presence of food, which conflicts with our findings but may result from differences in experimental conditions. In our experimental paradigm, animals were all adapted to the experimental environment for 2h in the morning before CNO administration, a period during which they had no access to food or food-related sensory cues. We restored access to food immediately after CNO administration, whereas in the Krashes paper, food was present before and after the injection. This suggests that changes in environmental food availability via food presentation are necessary to inhibit the AGRP-iBAT circuit, and that environmental food cues do not maintain a tonic inhibition of AGRP neurons. This interpretation is in fact consistent with our observation that caged-food only transiently blunts AGRP-induced suppression of energy expenditure and iBAT temperature.

In a study using capsaicin-dependent chemogenetic activation of AGRP neurons, activation of AGRP neurons produced a modest and transient decrease in energy expenditure that was attributed to a rapid suppression of inguinal WAT (iWAT) browning^15^. These authors ruled out a role for iBAT in this effect based on the lack of change in iBAT thermogenic gene expression ^15^. This latter observation is consistent with ^28^, our results, and a reported half-life of 30-72h of UCP1 protein in iBAT ^29^. Nonetheless, our data indicate that iBAT temperature can rapidly decrease following AGRP neuronal activation through activation of sympathetic signaling to iBAT and decreased activation of HSL, an enzyme implicated in fatty acid mobilization and heat production from iBAT. We can not exclude that a decrease in UCP1 expression in iWAT may contribute to the rapid decrease in energy expenditure observed following AGRP neuronal activation. However, we report an average decrease in energy expenditure of 8 J/min at ambient temperature (25% decrease compare to controls and baseline levels). While iBAT thermogenesis can account for this portion of energy expenditure in mice housed at ambient temperature ^30^, current knowledge does not indicate that it is the case for beige adipocyte ^31^ further supporting a major role of iBAT inhibition in the decrease in energy expenditure induced by AGRP neuronal activation.

Recently, AGRP neurons have been implicated in the regulation of insulin sensitivity via iBAT myostatin expression ^28^. This work extends the orchestration of the metabolic adaptations to energy deficit by AGRP neurons to include a suppression of peripheral glucose uptake, which may serve as a glucose sparing process to protect the brain from hypoglycemia in conditions of low energy availability.

The neurochemistry and neuroanatomical circuits underpinning AGRP neuronal regulation of iBAT remain to be fully characterized. Circuits downstream from AGRP neurons previously implicated in the modulation iBAT thermogenesis include NPYergic ^32^, melanocortinergic ^7, 33-34^, and GABA-ergic circuits ^35^. This diversity may underlie the coexistence of multiple circuits engaged under different metabolic situations or within different time frames ^13^, to adjust iBAT thermogenic function in response to changes in AGRP tone. Consistent with the idea that multiple circuits may contribute to the AGRP-iBAT axis, AGRP projections to the LH and the anterior bed nucleus of the stria terminalis have both been shown to mediate the effect of AGRP neurons on iBAT myostatin expression ^28^. Interestingly, although the paraventricular nucleus of the hypothalamus (PVH) receives dense axonal input from AGRP neurons^36^ and has been implicated in the regulation of sympathetic output to iBAT ^37-38^, AGRP projections to the PVH do not modulate iBAT myostatin expression. While a large proportion of PVH neurons negatively regulate iBAT thermogenesis ^37,39^, PVH oxytocin neurons positively regulate sympathetic tone to iBAT ^38^ and have been implicated in diet-induced thermogenesis ^40^. However, whether AGRP neurons project to PVH oxytocin neurons is not clear ^41,42^. PVH TH neurons represent an alternative population that may be involved in the AGRP-iBAT axis, as tonic inhibition from ARH NPY neurons to PVH TH neurons via Y1R signaling has also recently been implicated in the control of iBAT thermogenesis ^32^. DMH neurons may alternatively or concomitantly mediate the effect of AGRP neuronal activation on iBAT thermogenesis, as the DMH is a major source of sympathoexcitatory input to medullary iBAT sympathetic premotor neurons in rRPa ^38^, and GABAergic inhibitory tone to the DMH suppresses iBAT thermogenesis ^43^, raising the possibility that iBAT-regulating GABAergic input to the DMH originates from AGRP neurons.

Ghrelin is a well-established endogenous activator of AGRP neurons and the ghrelin receptor is predominantly expressed in AGRP neurons in the ARH ^44^. Our observation that parenchymal administration of ghrelin in the ARH reproduces the effects of AGRP neuronal activation on iBAT temperature supports the physiological relevance of the AGRP-iBAT circuit. While the contribution of iBAT thermogenesis to the metabolic effects of ghrelin has been previously reported ^45-46^, the neuroanatomical sites responsible for this effect had not been identified. Together with the published literature implicating AGRP neurons in the orexigenic effects of ghrelin ^47^,^48^, our data lend strength to the idea that AGRP neurons coordinate the feeding and metabolic effects of ghrelin.

Our findings establish a role for mTORC1 signaling in AGRP neurons in the detection of sensory and metabolic signals of energy availability and the regulation of the AGRP-iBAT axis. These findings may seem conflicting with a previous report showing increased ribosomal protein S6 (rpS6) signaling in AGRP neurons of fasted mice ^49^. In fact, rpS6 signaling receives activatory inputs from many neuronal signaling pathways, is now acknowledged as a marker of neuronal activation ^50^ and therefore represents an inappropriate surrogate marker of mTORC1 signaling in this context. Our data support instead that mTORC1 activation in AGRP neurons negatively regulates AGRP activity. Furthermore, we provide evidence that increased mTORC1 signaling in AGRP neurons contributes to diet-induced thermogenesis during early exposure to HF feeding and protects against diet-induced obesity (DIO). Consistent with these findings, acute HF feeding has been shown to increase S6K1 activity ^51^ and reduce AGRP signaling ^52^.

While there is convincing evidence that acute and chronic energy excess result in increased energy expenditure in rodents ^22^ and humans ^53,54^, the nature of the signals mediating these effects is unclear and may include specific nutrients and/or caloric load per se. Interestingly, the lack of effect of increased mTORC1 signaling in AGRP neurons under chow maintenance reveals that activation of this pathway is not sufficient to modulate energy balance, and suggests that its contribution to energy expenditure is contingent on energy surfeit. Controversy persists on the role of leptin during energy surfeit in diet-induced hypermetabolism humans ^55,56^, and recent data revealed novel mechanisms through which leptin modulates core temperature independently of energy expenditure ^57^. Nevertheless, in rodents, leptin increases sympathetic output to iBAT, oxygen consumption and iBAT temperature ^58 59 24^, and neuroanatomical and functional studies indicate that activation of central leptinergic circuits can rapidly increase iBAT thermogenesis ^60^. Our study identifies AGRP neurons as a novel neurochemical population implicated in leptin’s thermogenic action, although they do not directly implicate iBAT thermogenesis in the thermogenic response to leptin.

Previous work investigating the role of hypothalamic mTORC1 signaling support the prediction that modulation of S6K1 in AGRP neurons would affect food intake ^18,27^. The failure to observe any effect on feeding in our different experimental paradigms indicates that chronic changes in mTORC1 signaling in AGRP neurons are not sufficient to alter energy intake. This does not preclude a contribution of this pathway in the acute control of feeding behavior. In fact, we observe that rapamycin rapidly activates AGRP neurons in refed mice, and under similar conditions hypothalamic rapamycin has been shown to rapidly increase foraging and food intake ^18^. Here we demonstrate that rapamycin selectively suppresses iBAT temperature and energy expenditure in the absence of food, precisely mirroring the metabolic effects of AGRP neuronal activation. Interestingly, we observed that only a portion of AGRP neurons express active mTORC1 following a refeed or 2 days of HF feeding, indicating that the mTORC1 pathway is nutritionally regulated selectively in a subset of this neuronal population. Our findings contrast with other models of altered mTORC1 signaling in AGRP neurons, in which neonatal deletion of S6K1 or raptor in AGRP neurons failed to affect energy balance ^61,62^. Importantly, in contrast to the aforementioned studies, our model identifies the role of mTORC1 signaling in AGRP neurons in the physiology of mice with normally-developed hypothalamic circuits.

All together, our studies uncover a role for mTORC1 signaling within AGRP neurons in surveying energy availability to engage iBAT thermogenesis and identify AGRP neurons as a neuronal substrate for the coordination of energy intake and adaptive expenditure under various physiological contexts.

## Methods

### Animals

All mice were group-housed unless otherwise stated and maintained in individually ventilated cages with standard bedding and enrichment. Mice were maintained in a temperature and humidity-controlled room on a 12-hour light/dark cycle with *ad libitum* access to water and standard laboratory chow diet or HF diet D12266B, 31.8%k kcal fat, 4.4 kcal/g, Research Diets) unless otherwise stated. *Agrp-Ires-cre* mice were genotyped with the following primers: 52370- GGGCCCTAAGTTGAGTTTTCCT-32370; 52370-GATTACCCAACCTGGGCAGAAC-32370 and 52370- GGGTCGCTACAGACGTTGTTTG-32370. All experiments were carried on heterozygous *Agrp-IRES-cre* mice and their wild-type littermates. *NPY-GFP* mice were genotyped with the following primers: 5’- TATGTGGACGGGGCAGAAGATCCAGG-3’; 5’-CCCAGCTCACAT ATTTATCTAGAG-3’; 5’- GGTGCGGTTGCCGTACTGGA-3’. *NPY-GFP/Agrp-Ires-Cre* mice were obtained by crossing homozygous NPY-GFP mice with homozygous Agrp-Ires-Cre mice. All experiments were performed in accordance with the Animals (Scientific Procedures) Act 1986 and approved by the local animal ethic committees.

### Surgical procedures

Surgical procedures were performed under isofluorane anesthesia, and all animals received Metacam prior to the surgery, 24h after surgery and were allowed a 1-week recovery period during which they were acclimatized to injection procedures. Mice were stereotactically implanted with bilateral steel guide cannulae (Plastics One) positioned 1mm above the ARH (A/P: −1.1mm, D/V: −4.9mm, lateral: +0.4mm from Bregma), as previously described ^63^. Beveled stainless steel injectors (33 gauge) extending 1mm from the tip of the guide were used for injections. All viral brain injections were performed on 11-week old mice at 100nl/min, 500nl/side (lentivectors, 1–2 × 10^9^ pfu/ml) and 200 nl/side (AAV-hSyn-DIO-hM3D(Gq)-mCherry, UNC Vector Core, 1×10^12^ pfu/ml). For chronic cannulae implantation, cannula guide were secured in place with Loctite glue and dental cement (Fujicem). Correct targeting was confirmed histologically postmortem (placement of cannula guide track) or using mCherry immunofluorescent staining. Temperature telemetric probe (IPTT-300, BMDS) were inserted subcutaneously or secured below the interscapular brown adipose pad. All subsequent functional studies were performed in a crossover randomized manner on age- and weight-matched groups after 1-week recovery and 1-week daily acclimatization, and at least 4 days elapsed between each injection.

### Preparation of lentiviral plasmids

pRK7-HA-F5A (Addgene 8986) and pRK7-HA-KR (Addgene 8985) were a gift from John Blenis ^64^. pRK7-HA-F5A and pRK7-HA-KR were digested with XbaI followed by a fill-in reaction for blunt end ligation and a digestion with EcoRI to isolate the HA-F5A and HA-KR constructs. A lentiviral construct previously developed to express hTXNIP in a cre-dependent manner ^63^, pCDH-CMV-FLEX-hTXNIP, was digested with Xho-1 followed by a fill-in reaction for blunt end ligation and a digestion with EcoR1 to isolate the pCDH-CMV-FLEX vector. The 2 inserts were ligated to the pCDH-CMV-FLEX lentivector to produce pCDH-CMV-FLEX-Ha-S6K1-F5A and pCDH-CMV-FLEX-Ha-S6K1-KR. Several clones were screened using One Shot Stbl3 E Coli competent cells (ThermoFisher and verified by sequencing. Plasmids were subsequently used for packaging reactions to generate viral stocks suitable for transfection by System Bioscience (Mountain View, CA). Cre-dependent expression of S6K1 constructs was confirmed in HEK293T cells (ATCC, negative for mycoplasma contamination) transfected with a pCAG-Cre-IRES2-GFP plasmid, a gift from Dr. Anjen Chenn’s lab (Addgene 26646).

### Metabolic phenotyping

#### AGRP chemogenetic activation

On the day of the procedure, mice housed in their home cage were acclimatized to the procedure room and food deprived during the 2h preceding the injections. Mice received CNO (Sigma, 1mg/kg) or vehicle and were maintained in a food-free cage for the four following hours unless otherwise stated. iBAT or subcutaneous temperature were measured using remote biotelemetry (IPTT-300 Bio Medic Data Systems). Energy expenditure was assessed using indirect calorimetry in a Metabolic-Trace system (Meta-Trace, Ideas Studio, UK).

#### Brain injections

For ghrelin or leucine injection studies, mice housed in their home cage were acclimatized to the procedure room and food deprived during the 2h preceding the injections. Mice received bilateral parenchymal nanoinjection of ghrelin (Phoenix Pharmaceuticals, 1mg/ml, 100nl/side, 100nl/min), L-leucine (Sigma, 2.1mM, 100nl/side, 100nl/min) or aCSF (R&D), were immediately returned to their home cage, and iBAT temperature and energy expenditure were monitored as described above. For rapamycin injection studies, mice were fasted overnight, re-fed for 1h in the morning, nanoinjected with rapamycin or DMSO (Merck Millipore, 5mM, 150nl/side), placed in clean, food-free cages, and iBAT temperature and energy expenditure were monitored as described above.

#### Oral Glucose Tolerance

Glucose tolerance was measured on body-weight matched animals. Mice were food deprived for 6h and blood was sampled from tail vein immediately prior to glucose bolus (gavage, 1mg/kg), and 15, 30, 60 and 120 minutes following bolus administration. Blood glucose was analyzed using a glucometer (Precision Xtra; MediSense).

#### Body Composition Analysis

Body composition was determined by magnetic resonance spectroscopy (MRS) using an Echo MRS instrument (Echo Medical Systems).

#### CL challenge

After blood collection for basal measurements, mice received an intraperitoneal injection of 1 mg/kg BW CL316243 (5-[(2*R*)-2-[[(2*R*)-2-(3-chlorophenyl)-2-hydroxyethyl]amino]propyl]-1,3-benzodioxole-2,2-dicarboxylic acid disodium salt; Sigma), a β3 adrenergic agonist. Brown fat temperature was monitored as described above, over the 60 min after the CL316243 injection, and brown fat temperature change over that time period was calculated. *Leptin challenge*. Mice received an intraperitoneal injection of leptin (5mg/kg, R&D) or saline 1h before the onset of the dark, and iBAT temperature, 24h body weight gain and 24h food intake were measured.

### RT-qPCR

qPCR was carried out as previously described ^65^. Total RNA was purified from interscapular iBAT using RNA STAT 60 (AMS Biotechnology, Abington, UK) according to the manufacturer’s instructions. cDNA was obtained by reverse transcription of 500 ng iBAT RNA. PCR of cDNA was performed in duplicate on an ABI Prism 7900 sequence detection system using Taqman Gene expression assay for *Elovl3, Pgc-1a, Ucp-1 and Dio2* (Supplement 1). Data expressed as arbitrary units and expression of target genes corrected to the geometric average of four housekeeping genes: *18s, 36β4, βactin* and *Gapdh*.

### Western Blots

Tissues were ground to a fine powder using a sterile pestle and mortar on liquid nitrogen. Powdered tissue were resuspended in lysis buffer (50 mM Tris-HCL, 150 mM NaCl, 1 mM EGTA, 1 mM EDTA, 10mM glycerophosphate, 2mM orthovanadate, 2mM PMSF, .5% sodium deoxycholate, 1% Triton X-100, pH7.5) with added protease and phosphatase inhibitor cocktails according to manufacturers instruction (Roche). Lysates were cleared by centrifugation at 2600g for 5 min at 4 °C, to separate the fat content of the samples. Protein extracts were further cleared by centrifugation at 10 000*g* for 10 min at 4 °C. Protein concentrations of the supernatants were determined using the Bradford assay, proteins were diluted in Laemli buffer and separated by SDS–polyacrylamide gel (12%) electrophoresis and transferred to Immobilon-P (Millipore) membrane). Membranes were blocked for 12370h at room temperature and incubated overnight at 4 °C with the indicated antibody (1:1000, Cell Signaling Technology). Bound primary antibodies were detected using peroxidase-coupled secondary antibodies and enhanced chemiluminescence (Amersham). Relative quantification of band intensities was calculated by digitally photographing exposed films and using Genesnap and Genetools software (Syngene).

### Brain perfusion, immunohistochemistry, confocal microscopy and image analysis

Animals were anaesthetized with Euthatal solution 80 mg/kg in saline and transcardiacally perfused with 4% paraformaldehyde. Brains were extracted and post-fixed in 4% paraformaldehyde, 30% sucrose for 48h at 4 °C. Brains were sectioned using a freezing sliding microtome into 5 subsets of 25 microns sections. Antigen retrieval was used for all experiments prior to antibody incubation. Sections were incubated in 10 mM sodium citrate at 80? for 20min then washed 3 times in PBS. To decrease endogenous autofluorescence, samples were then treated with 0.3% glycine and 0.003% SDS in methanol. Tissue was blocked for 1h with 5% normal donkey serum at room temperature, and incubated at 4 °C with primary antibodies against pSer2448-mTORC1 (72h, 1:65; Cell Signaling Technology), c-fos (48h, 1:5000, Synaptic Systems), pSer240/244rpS6 (19h, 1:200, Cell Signaling Technology), dsRed (1:1000, Clontech), or pSTAT3 (19h, 1:500, Cell Signaling Technology). Sections were then mounted on slides and coverslipped with Vectashield (Vector).

Sections were imaged using a Zeiss LSM510 confocal microscope with the 40x objective. Gain and laser power settings remained the same between experimental and control conditions. Images of tissue sections were digitized, and areas of interest were outlined based on cellular morphology. The brain region evaluated was arcuate nucleus (1.5–1.9 mm caudal, 0–0.45 mm mediolateral, and 5.6–6 mm ventromedial to bregma), corresponding to the coordinates in the brain atlas of ^21^. Images were analyzed using the ImageJ manual cell counter.

### Serum analysis

Mouse FGF-21 was measured using a Quantikine ELISA kit from Bio-Techne (MF2100). The assay was performed according to the manufacturer’s instructions. Samples were analyzed in duplicate and the mean value reported. Quality control samples were analyzed at the beginning and end of the batch to ensure consistency across the plate.

T4 was measured using a fully automated assay on the Siemens Dimension EXL analyzer. The method is an adaptation of the EMIT homogenous immunoassay technology. T4 is stripped from its binding proteins using a releasing agent. The released T4 binds to anti-T4 antibody, thereby reducing the amount of T4-glucose-6-phosphate dehydrogenase conjugate (T4-G6PDH) that can be bound to the antibody. The T4-G6PDH which is not bound to the antibody catalyzes the oxidation of glucose-6-phosphate with the simultaneous reduction of NAD^+^ to NADH which absorbs at 340 nm. In the absence of T4, T4-G6PDH binds with the antibody and enzyme activity is reduced. The increase in absorbance at 340 nm due to the formation of NADH over a 60 second measure period is proportional to the activity of the T4-G6PDH. The T4 concentration is measured using a bichromatic (340, 383 nm) rate technique. All reagents and calibrators were purchased from Siemens. Samples were analyzed in singleton. Three levels of Quality Control samples were analyzed at the beginning and end of the batch. All QC results were within their target ranges before the sample results were reported.

### iBAT Norepinephrine turnover

iBAT norepinephrine turnover was determined based on the fall in iBAT norepinephrine content following inhibition of catecholamine biosynthesis with α-methyl-DL-tyrosine methyl ester hydrochloride (AMPT,Sigma-Aldrich) administered ip at 250 mg/kg, as described previously ^66^. AMPT is a competitive inhibitor of tyrosine hydroxylase, the rate-limiting enzyme in catecholamine biosynthesis. After AMPT administration, the endogenous tissue levels of norepinephrine decline and the slope of this decrease in tissue norepinephrine content is multiplied by the initial norepinephrine concentration to yield NETO, an indirect readout of SNS outflow.

Norepinephrine was assayed in iBAT using reversed phase high performance liquid chromatography (HPLC) and electrochemical detection ^67^. Samples (20mg) were homogenized in 0.2 ml of 0.2M perchloric acid and centrifuged at 8000 rpm for 20 min at 10oC. The supernatant was diluted 1:4 in PCA buffer before the injection. Twenty-five μl aliquots of supernatant were injected onto a C18 ODS 3μm column (100 mm length x 4.6 mm i.d.,Hypersil Elite, Phenomenex, UK) with a mobile phase consisting of citric acid (31.9 g/L), sodium acetate (2 g/L), octanesulfonic acid (460 mg/L) EDTA (30 mg/L) and 15% methanol (pH 3.6). Norepinephrine was detected using an ESA Coulochem II detector with electrode 1 held at −200 mV and electrode 2 held at +250 mV. Chromatograms were acquired and analyzed using Chromeleon software (Dionex, UK).

### Statistical analysis

All data, presented as means ± SEM, have been analyzed using GraphPad Prism 6. For all statistical tests, an α risk of 5% was used. All kinetics were analyzed using repeated-measures two-way ANOVAs and adjusted with Bonferroni’s post hoc tests. Multiple comparisons were tested with one-way ANOVAs and adjusted with Tukey’s post hoc tests. Single comparisons were made using one-tail Student’s t tests.

## Acknowledgment

This work was supported by the Medical Research Council New Blood Fellowship [MR/M501736/1] to CB, a NIDDK K99/R00 award to CB, the Medical Research Council Metabolic Disease Unit programme grant, the Disease Model Core facilities, and the Wellcome Trust Cambridge Mouse Biochemistry Laboratory. Animal procedures were performed under Tony Coll’s home office PPL 80/2497 and Toni Vidal-Puig PPL 80/2484.

## Figure Supplements Legend

**Figure 1-Supplement 1.** Confirmation of successful AAV-targeting using mCherry staining - Scale bar 150 μm (a). Confirmation of increased food intake upon CNO administration (b). iBAT temperature in response to local nanoinjection of ghrelin or artificial cerebrospinal fluid (aCSF) into the mediobasal hypothalamuc (MBH) (c). Serum thyroxine (d) and FGF21 (e) concentrations before and 45 min after AGRP activation. Thermogenic gene expression (f) and β3-adrenergic signaling (g) in iBAT 2h following AGRP activation. iBAT temperature response to AGRP neuronal activation in animals maintained a 4°C (h). Data are mean ± SEM. ***: p<0.001

**Figure 2-Supplement 1.** iBAT temperature (a) and energy expenditure (b) following chemogenetic activation of AGRP neurons. iBAT temperature (a) and energy expenditure (b) following chemogenetic activation of AGRP neurons and exposure to a novel empty container. Data are mean ± SEM.

**Figure 3-Supplement 1.** iBAT temperature (a) and energy expenditure (b) in mice fasted for 19h and refed for 60 min. p-mTOR immunofluorescent staining in mouse hypothalamic brain sections at the rostrocaudal level of the ARH 60min following an ip injection of DMSO or rapamycin (c).p-mTOR (red) immunostaining in 1 refed NPY-GFP mouse showing the rostrocaudal localization of the NPY/p-mTOR costain (d). iBAT temperature (e) iBAT *ucp-1* mRNA expression (f) and energy expenditure (g) in mice maintained on chow or fed a high fat diet for 48h. Locomotor activity (h) in mice fasted overnight for 19h, refed a chow meal and injected with rapamycin into the MBH. Energy expenditure (i) in mice fasted for 6h and injected with L-leucine into the MBH. Data are mean ± SEM. *: p<0.05, ***: p<0.001

**Figure 4-Supplement 1.** Ha (a) and p-rpS6 (b) immunostaining in Agrp-IRES-cre;Npy-gfp mice infected with the pCDH-CMV-FLEX-HA-S6K1-F5a lentivirus. Body weight gain (c) fat mass (d) and glucose tolerance (d) of Agrp-CA-S6 and WT-CA-S6 mice under chow maintenance. Data are mean ± SEM

**Figure 5-Supplement 1.** Ha (a) and p-rpS6 (b) immunostaining in Agrp-IRES-cre;Npy-gfp mice infected with the pCDH-CMV-FLEX-HA-S6K1-KR lentivirus. Body weight gain on chow (c) and RER in Agrp-DN-S6 and WT-DN-S6 mice. Data are mean ± SEM

**Supplementary file 1.** Real time PCR Oligonucleotide Sequences

## REFERENCES

1 Bachman, E. S. et al. betaAR signaling required for diet-induced thermogenesis and obesity resistance. Science (New York, N.Y). 297, 843–845 (2002).

2 Feldmann, H. M., Golozoubova, V., Cannon, B. & Nedergaard, J. UCP1 ablation induces obesity and abolishes diet-induced thermogenesis in mice exempt from thermal stress by living at thermoneutrality. Cell metabolism 9, 203–209, doi:10.1016/j.cmet.2008.12.014 (2009).

3 Ravussin, Y., LeDuc, C. A., Watanabe, K. & Leibel, R. L. Effects of ambient temperature on adaptive thermogenesis during maintenance of reduced body weight in mice. American journal of physiology. Regulatory, integrative and comparative physiology 303, R438–448, doi:10.1152/ajpregu.00092.2012 (2012).

4 Leibel, R. L., Rosenbaum, M. & Hirsch, J. Changes in energy expenditure resulting from altered body weight. The New England journal of medicine 332, 621–628, doi:10.1056/nejm199503093321001 (1995).

5 Bartness, T. J., Shrestha, Y. B., Vaughan, C. H., Schwartz, G. J. & Song, C. K. Sensory and sympathetic nervous system control of white adipose tissue lipolysis. Mol Cell Endocrinol (2009).

6 Brito, N. A., Brito, M. N. & Bartness, T. J. Differential sympathetic drive to adipose tissues after food deprivation, cold exposure or glucoprivation. Am J Physiol Regul Integr Comp Physiol. 294, R1445–1452. Epub 2008 Mar 1445. (2008).

7 Small, C. J. et al. Chronic CNS administration of Agouti-related protein (Agrp) reduces energy expenditure. International journal of obesity and related metabolic disorders: journal of the International Association for the Study of Obesity 27, 530–533, doi:10.1038/sj.ijo.0802253 (2003).

8 Chen, Y., Lin, Y. C., Kuo, T. W. & Knight, Z. A. Sensory detection of food rapidly modulates arcuate feeding circuits. Cell 160, 829–841, doi:10.1016/j.cell.2015.01.033 (2015).

9 Konner, A. C. et al.. Insulin action in AgRP-expressing neurons is required for suppression of hepatic glucose production. Cell Metab. 5, 438–449. (2007).

10 Luquet, S., Perez, F. A., Hnasko, T. S. & Palmiter, R. D. NPY/AgRP neurons are essential for feeding in adult mice but can be ablated in neonates. Science. 310, 683–685. (2005).

11 Gropp, E. et al.. Agouti-related peptide-expressing neurons are mandatory for feeding. Nat Neurosci. 8, 1289–1291. Epub 2005 Sep 1211. (2005).

12 Aponte, Y., Atasoy, D. & Sternson, S. M. in Nature neuroscience Vol. 14 351–355 (2011).

13 Krashes, M. J. et al.. Rapid, reversible activation of AgRP neurons drives feeding behavior in mice. The Journal of clinical investigation 121, 1424–1428, doi:10.1172/jci46229 (2011).

14 Alexander, G. M. et al.. Remote control of neuronal activity in transgenic mice expressing evolved G protein-coupled receptors. Neuron 63, 27–39, doi:10.1016/j.neuron.2009.06.014 (2009).

15 Ruan, H. B. et al.. O-GlcNAc transferase enables AgRP neurons to suppress browning of white fat. Cell 159, 306–317, doi:10.1016/j.cell.2014.09.010 (2014).

16 Anthony, N. M., Gaidhu, M. P. & Ceddia, R. B. Regulation of visceral and subcutaneous adipocyte lipolysis by acute AICAR-induced AMPK activation. Obesity (Silver Spring, Md.) 17, 1312–1317, doi:10.1038/oby.2008.645 (2009).

17 Pulinilkunnil, T. et al.. Adrenergic regulation of AMP-activated protein kinase in brown adipose tissue in vivo. The Journal of biological chemistry 286, 8798–8809, doi:10.1074/jbc.M111.218719 (2011).

18 Cota, D. et al.. Hypothalamic mTOR signaling regulates food intake. Science (New York, N.Y). 312, 927–930, doi:312/5775/927 [pii]10.1126/science.1124147 [doi] (2006).

19 Blouet, C. & Schwartz, G. J. Brainstem nutrient sensing in the nucleus of the solitary tract inhibits feeding. Cell metabolism 16, 579–587 (2012).

20 Teubner, B. J., Keen-Rhinehart, E. & Bartness, T. J. Third ventricular coinjection of subthreshold doses of NPY and AgRP stimulate food hoarding and intake and neural activation. American journal of physiology. Regulatory, integrative and comparative physiology 302, R37–48, doi:10.1152/ajpregu.00475.2011 (2012).

21 Paxinos, G. & Franklin, K. B. J. The Mouse Brain in Stereotaxic Coordinates, Second edition. Academic Press, NY (2001).

22 Cannon, B. & Nedergaard, J. Brown adipose tissue: function and physiological significance. Physiol Rev. 84, 277–359. (2004).

23 Liu, C. et al.. PPARgamma in vagal neurons regulates high-fat diet induced thermogenesis. Cell metabolism 19, 722–730, doi:10.1016/j.cmet.2014.01.021 (2014).

24 Rahmouni, K. & Morgan, D. A. Hypothalamic arcuate nucleus mediates the sympathetic and arterial pressure responses to leptin. Hypertension (Dallas, Tex.: 1979) 49, 647–652, doi:01.HYP.0000254827.59792.b2 [pii]10.1161/01.HYP.0000254827.59792.b2 [doi] (2007).

25 Harlan, S. M. et al.. Ablation of the leptin receptor in the hypothalamic arcuate nucleus abrogates leptin-induced sympathetic activation. Circulation research 108, 808–812, doi:10.1161/circresaha.111.240226 (2011).

26 Baquero, A. F. et al.. Developmental switch of leptin signaling in arcuate nucleus neurons. The Journal of neuroscience: the official journal of the Society for Neuroscience 34, 9982–9994, doi:10.1523/jneurosci.0933-14.2014 (2014).

27 Blouet, C., Ono, H. & Schwartz, G. J. Mediobasal hypothalamic p70 S6 kinase 1 modulates the control of energy homeostasis. Cell Metab. 8, 459–467. (2008).

28 Steculorum, S. M. et al.. AgRP Neurons Control Systemic Insulin Sensitivity via Myostatin Expression in Brown Adipose Tissue. Cell 165, 125–138, doi:10.1016/j.cell.2016.02.044 (2016).

29 Clarke, K. J. et al.. A role for ubiquitinylation and the cytosolic proteasome in turnover of mitochondrial uncoupling protein 1 (UCP1). Biochimica et biophysica acta 1817, 1759–1767, doi:10.1016/j.bbabio.2012.03.035 (2012).

30 Cannon, B. & Nedergaard, J. Nonshivering thermogenesis and its adequate measurement in metabolic studies. The Journal of experimental biology 214, 242–253, doi:10.1242/jeb.050989 (2011).

31 Nedergaard, J. & Cannon, B. UCP1 mRNA does not produce heat. Biochimica et biophysica acta 1831, 943–949, doi:10.1016/j.bbalip.2013.01.009 (2013).

32 Shi, Y. C. et al.. Arcuate NPY controls sympathetic output and BAT function via a relay of tyrosine hydroxylase neurons in the PVN. Cell metabolism 17, 236–248, doi:10.1016/j.cmet.2013.01.006 (2013).

33 Berglund, E. D. et al.. Melanocortin 4 receptors in autonomic neurons regulate thermogenesis and glycemia. Nature neuroscience 17, 911–913, doi:10.1038/nn.3737 (2014).

34 Brito, M. N., Brito, N. A., Baro, D. J., Song, C. K. & Bartness, T. J. Differential activation of the sympathetic innervation of adipose tissues by melanocortin receptor stimulation. Endocrinology. 148, 5339–5347. Epub 2007 Aug 5316. (2007).

35 Tong, Q., Ye, C. P., Jones, J. E., Elmquist, J. K. & Lowell, B. B. Synaptic release of GABA by AgRP neurons is required for normal regulation of energy balance. Nat Neurosci. 11, 998–1000. (2008).

36 Haskell-Luevano, C. et al.. Characterization of the neuroanatomical distribution of agouti-related protein immunoreactivity in the rhesus monkey and the rat. Endocrinology 140, 1408–1415, doi:10.1210/endo.140.3.6544 (1999).

37 Kong, D. et al.. GABAergic RIP-Cre neurons in the arcuate nucleus selectively regulate energy expenditure. Cell 151, 645–657, doi:10.1016/j.cell.2012.09.020 (2012).

38 Morrison, S. F. & Madden, C. J. Central nervous system regulation of brown adipose tissue. Comprehensive Physiology 4, 1677–1713, doi:10.1002/cphy.c140013 (2014).

39 Madden, C. J. & Morrison, S. F. Neurons in the paraventricular nucleus of the hypothalamus inhibit sympathetic outflow to brown adipose tissue. American journal of physiology. Regulatory, integrative and comparative physiology 296, R831–843, doi:10.1152/ajpregu.91007.2008 (2009).

40 Wu, Z. et al.. in PloS one Vol. 7 e45167 (2012).

41 Atasoy, D. et al.. in Nature neuroscience Vol. 17 1830–1839 (2012).

42 Garfield, A. S. et al.. A neural basis for melanocortin-4 receptor-regulated appetite. Nature neuroscience 18, 863–871, doi:10.1038/nn.4011 (2015).

43 Cao, W. H., Fan, W. & Morrison, S. F. Medullary pathways mediating specific sympathetic responses to activation of dorsomedial hypothalamus. Neuroscience 126, 229–240, doi:10.1016/j.neuroscience.2004.03.013 (2004).

44 Willesen, M. G., Kristensen, P. & Romer, J. Co-localization of growth hormone secretagogue receptor and NPY mRNA in the arcuate nucleus of the rat. Neuroendocrinology 70, 306–316 (1999).

45 Tsubone, T. et al.. Ghrelin regulates adiposity in white adipose tissue and UCP1 mRNA expression in brown adipose tissue in mice. Regulatory peptides 130, 97–103, doi:10.1016/j.regpep.2005.04.004 (2005).

46 Yasuda, T., Masaki, T., Kakuma, T. & Yoshimatsu, H. Centrally administered ghrelin suppresses sympathetic nerve activity in brown adipose tissue of rats. Neuroscience letters 349, 75–78 (2003).

47 Wang, Q. et al.. Arcuate AgRP neurons mediate orexigenic and glucoregulatory actions of ghrelin. Molecular metabolism 3, 64–72, doi:10.1016/j.molmet.2013.10.001 (2014).

48 Andrews, Z. B. et al.. UCP2 mediates ghrelin’s action on NPY/AgRP neurons by lowering free radicals. Nature. 454, 846–851. Epub 2008 Jul 2030. (2008).

49 Villanueva, E. C. et al.. Complex regulation of mammalian target of rapamycin complex 1 in the basomedial hypothalamus by leptin and nutritional status. Endocrinology. 150, 4541–4551. (2009).

50 Knight, Z. A. et al.. Molecular profiling of activated neurons by phosphorylated ribosome capture. Cell 151, 1126–1137, doi:10.1016/j.cell.2012.10.03910.1016/j.cell.2012.10.039 (2012).

51 Cota, D., Matter, E. K., Woods, S. C. & Seeley, R. J. The role of hypothalamic mammalian target of rapamycin complex 1 signaling in diet-induced obesity. J Neurosci. 28, 7202–7208. (2008).

52 Ziotopoulou, M., Mantzoros, C. S., Hileman, S. M. & Flier, J. S. Differential expression of hypothalamic neuropeptides in the early phase of diet-induced obesity in mice. American journal of physiology. Endocrinology and metabolism 279, E838–845 (2000).

53 Ravussin, Y., Leibel, R. L. & Ferrante, A. W., Jr. A missing link in body weight homeostasis: the catabolic signal of the overfed state. Cell metabolism 20, 565–572, doi:10.1016/j.cmet.2014.09.002 (2014).

54 Vosselman, M. J. et al.. Brown adipose tissue activity after a high-calorie meal in humans. The American journal of clinical nutrition 98, 57–64, doi:10.3945/ajcn.113.059022 (2013).

55 Heymsfield, S. B. et al.. Recombinant leptin for weight loss in obese and lean adults: a randomized, controlled, dose-escalation trial. Jama 282, 1568–1575 (1999).

56 Mackintosh, R. M. & Hirsch, J. The effects of leptin administration in non-obese human subjects. Obesity research 9, 462–469, doi:10.1038/oby.2001.60 (2001).

57 Fischer, A. W. et al.. Leptin Raises Defended Body Temperature without Activating Thermogenesis. Cell reports 14, 1621–1631, doi:10.1016/j.celrep.2016.01.041 (2016).

58 Morrison, S. F. Activation of 5-HT1A receptors in raphe pallidus inhibits leptin-evoked increases in brown adipose tissue thermogenesis. American journal of physiology. Regulatory, integrative and comparative physiology 286, R832–837, doi:10.1152/ajpregu.00678.2003 (2004).

59 Enriori, P. J., Sinnayah, P., Simonds, S. E., Garcia Rudaz, C. & Cowley, M. A. Leptin action in the dorsomedial hypothalamus increases sympathetic tone to brown adipose tissue in spite of systemic leptin resistance. The Journal of neuroscience: the official journal of the Society for Neuroscience 31, 12189–12197, doi:10.1523/jneurosci.2336-11.2011 (2011).

60 Yu, S. et al.. Glutamatergic Preoptic Area Neurons That Express Leptin Receptors Drive Temperature-Dependent Body Weight Homeostasis. The Journal of neuroscience: the official journal of the Society for Neuroscience 36, 5034–5046, doi:10.1523/jneurosci.0213- 16.2016 (2016).

61 Smith, M. A. et al.. Ribosomal S6K1 in POMC and AgRP Neurons Regulates Glucose Homeostasis but Not Feeding Behavior in Mice. Cell reports 11, 335–343, doi:10.1016/j.celrep.2015.03.029 (2015).

62 Albert, V., Cornu, M. & Hall, M. N. mTORC1 signaling in Agrp neurons mediates circadian expression of Agrp and NPY but is dispensable for regulation of feeding behavior. Biochemical and biophysical research communications 464, 480–486, doi:10.1016/j.bbrc.2015.06.161 (2015).

63 Blouet, C., Liu, S. M., Jo, Y. H., Chua, S. & Schwartz, G. J. in The Journal of neuroscience: the official journal of the Society for Neuroscience Vol. 32 9870–9877 (2012).

64 Schalm, S. S. & Blenis, J. Identification of a conserved motif required for mTOR signaling. Current biology: CB 12, 632–639 (2002).

65 Burke, L. K. & Heisler, L. K. 5-hydroxytryptamine medications for the treatment of obesity. Journal of neuroendocrinology 27, 389–398, doi:10.1111/jne.12287 (2015).

66 Brodie, B. B., Costa, E., Dlabac, A., Neff, N. H. & Smookler, H. H. Application of steady state kinetics to the estimation of synthesis rate and turnover time of tissue catecholamines. The Journal of pharmacology and experimental therapeutics 154, 493–498 (1966).

67 Dalley, J. W., Theobald, D. E., Eagle, D. M., Passetti, F. & Robbins, T. W. Deficits in impulse control associated with tonically-elevated serotonergic function in rat prefrontal cortex. Neuropsychopharmacology: official publication of the American College of Neuropsychopharmacology 26, 716–728, doi:10.1016/s0893-133x(01)00412-2 (2002).

